# Self-Assembled Monolayer Transporters Enable Reagentless Analysis of Small Molecule Analytes

**DOI:** 10.1101/2024.05.07.592958

**Authors:** Connor D. Flynn, Kimberly T. Riordan, Tiana L. Young, Dingran Chang, Zhenwei Wu, Scott E. Isaacson, Hanie Yousefi, Jagotamoy Das, Shana O. Kelley

## Abstract

The detection of small molecules beyond glucose remains an ongoing challenge in the field of biomolecular sensing owing to their small size, diverse structures, and lack of alternative non-enzymatic sensing methods. Here, we present a new reagentless electrochemical approach for small molecule detection that involves directed movement of electroactive analytes through a self-assembled monolayer to an electrode surface. Using this method, we demonstrate detection of several physiologically relevant small molecules as well as the capacity for the system to operate in several biological fluids. We anticipate that this mechanism will further improve our capacity for small molecule measurement and provide a new generalizable monolayer-based technique for electrochemical assessment of various electroactive analytes.

---

The detection of small molecules, including neurotransmitters, hormones, metabolites, and exogenous drugs, remains a key challenge in the pursuit of universal biomolecular sensing.^1,2^ While certain small molecule analytes (e.g., glucose, cholesterol) are easily detectable with electrochemical enzyme-based mechanisms, the scarcity of redox enzymes for other targets makes widespread adoption of this method difficult.^3^ Alternative sensing approaches including fast-scan cyclic voltammetry (FSCV) and electrochemical aptamer-based (E-AB) sensors offer promising solutions, but also come with their share of challenges; for example, FSCV frequently faces issues with sensitivity and selectivity, and E-AB sensors require aptamer recognition elements that undergo sufficient conformational changes upon small molecule binding.^4,5^ As such, the development of new techniques capable of sensing small molecule analytes remains an ongoing unmet need.

Recently, our lab published a new reagentless protein sensing approach using DNA constructs referred to as molecular pendulums (MPs).^6,7^ The MP sensing mechanism relies on potential-mediated movement of electrode-bound double-stranded DNA (dsDNA), containing both a terminal recognition element (i.e., antibody, aptamer) and terminal ferrocene molecule, toward an electrode surface.^8^ The rate of ferrocene oxidation near the electrode is modulated by the hydrodynamic radius of the MP, such that binding of a protein analyte slows MP descent and delays faradaic readout. Similar dynamic DNA modulation has also been demonstrated in optical systems employing DNA nanolevers.^9–11^ While MPs function well for proteins and larger analytes, their dependence on hydrodynamic radius changes makes them difficult to implement for smaller, less bulky targets. Here, we present a new approach that allows for reagentless analysis of electroactive small molecules. This method employs a dsDNA construct similar to the MP, with a terminal recognition element, but requires no external redox reporter; instead, the innate electroactivity of the small molecule analyte serves as the reporter, which is transported to the electrode surface using an applied potential (Figure 1a). We refer to these sensors as monolayer transporters (MTs) given their role in selectively binding and moving analytes of interest through the self-assembled monolayer to the electrode surface for oxidation.

**Figure 1.**
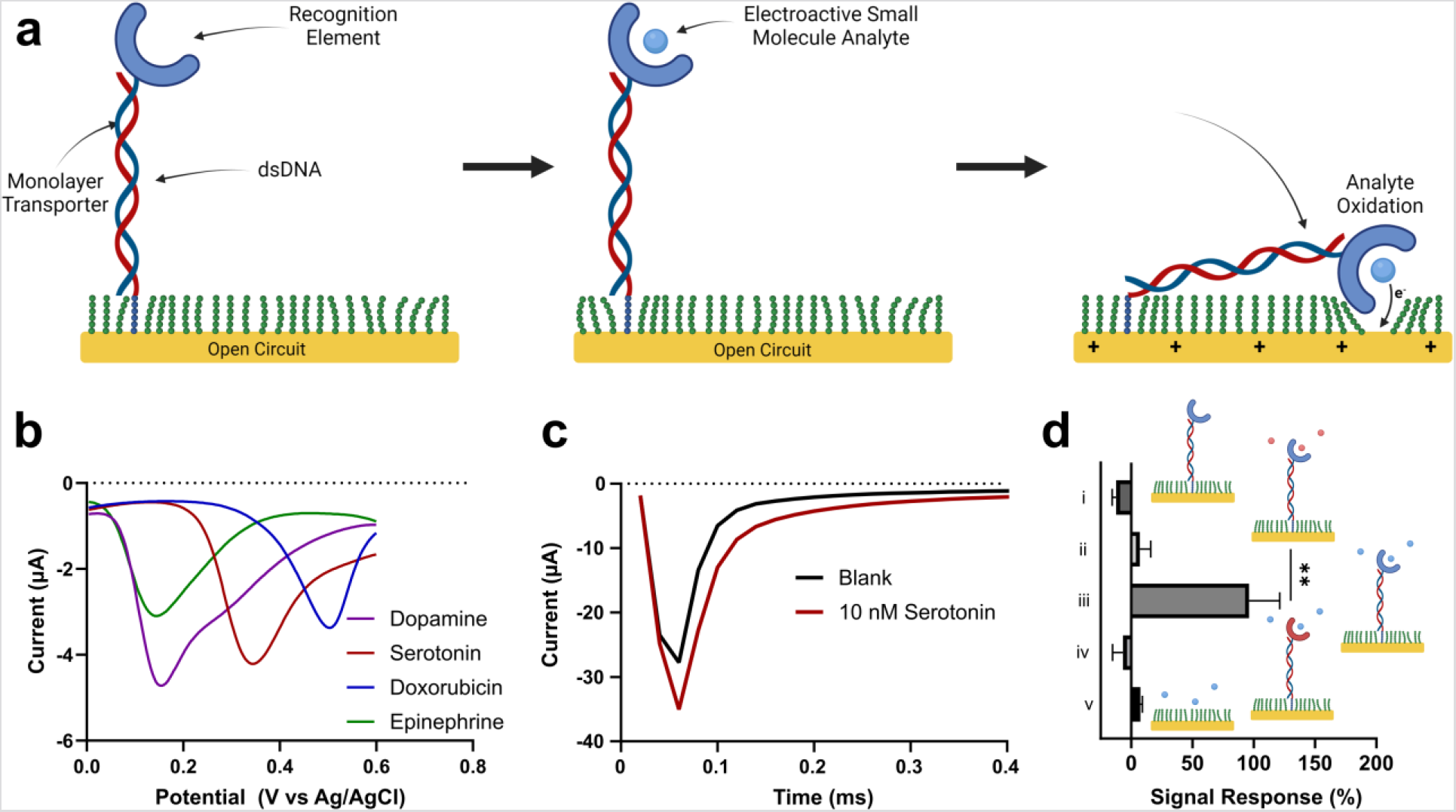
Monolayer Transporter (MT)-based Detection. (a) Schematic of MT-enabled detection of electroactive small molecules using a DNA-conjugated recognition element (e.g., aptamer, antibody). (b) Differential pulse voltammetry (DPV) measurements of the oxidation of several small molecules in PBS. (c) Chronoamperometric signal readout of serotonin-specific MTs before and after 30 min incubation in 10 nM serotonin. (d) Signal response reflected as percentages of serotonin-specific MTs after incubation in (i) PBS (ii) 10 µM dopamine (iii) 10 pM serotonin; (iv) dopamine-specific MTs after incubation in 10 µM serotonin; (v) MCH-only chip after incubation in 10 µM serotonin (30 min incubation and n = 5 for all).

A potential of +500 mV (vs. Ag/AgCl) is employed to bring the DNA construct to the electrode surface and facilitate oxidation of the analyte; this value represents a maximum as higher potentials can result in sensor degradation due to probe ejection or receptor breakdown.^6,12^ Within the current MT scheme, this restricts the small molecule targets that can be detected to those with an oxidation potential falling inside the 0-500 mV range. Fortunately, many biologically relevant small molecules reside within this range, owing to their structural similarities and common functional groups (e.g., phenols^13^). Figure 1b shows the oxidation of several small molecules that were explored in this study, with oxidation curves that fall at least partially within the 0-500 mV window. These selected analytes include both endogenous (e.g., dopamine) and exogenous (e.g., doxorubicin) targets. In addition to these molecules, many other neurotransmitters (e.g., norepinephrine^14^), hormones (e.g., melatonin^15^), lipids (e.g., cholesterol^16^), and drugs (e.g., acetaminophen^17^) have been reported with seemingly compatible oxidation potentials. MT sensors were prepared on cleanroom-fabricated gold microelectrodes containing additional electrodeposited gold nanostructures (Figure S1). These gold nanostructured microelectrodes (NMEs) were employed given prior reports of sensitivity enhancement^18^ and compatibility with previous MP-based sensing.^6,7^ Monolayers were prepared by sequential deposition of MT probes (hybridized anchor and receptor strands) and 6-merceptohexanol (MCH). Monolayer deposition was optimized to ensure stable chronoamperometric readout without compromising probe sensing ability. It should be noted that extensive MCH exposure led to minimal detection – either through probe replacement with MCH or through overly tight monolayers via thiol packing and reorganization. Likewise, too little MCH exposure led to overly porous monolayers that provided sufficient target signal but were also susceptible to non-specific oxidation events. Optimal probe density was determined to be approximately 4.3 ± 0.74 x 10^12^ probes/cm^2^ or 1 probe per 23 nm^2^ (Figure S3).

Using the optimized deposition protocol, serotonin-specific MTs were challenged with 10 nM serotonin in 1X phosphate-buffered saline (PBS), which produced a significant chronoamperometric response (Figure 1c). This response manifests as an increase in chronoamperometric current due to additional faradaic current from the oxidized analyte, which shifts the response waveform towards larger currents in the 40-300 µs window.

To further explore the viability of the monolayer transporter approach, as well as the integrity of the monolayer, several negative controls were tested in parallel alongside the positive target. Serotonin-specific MTs were assessed before and after incubation in PBS, a high concentration of negative target (10µM dopamine), and 10 pM serotonin. The negative controls showed negligible signal response (Figure 1d, i & ii), while serotonin exposure led to a noticeable response (Figure 1d, iii). Dopamine-specific MTs and MCH-only monolayers were also exposed to high concentration serotonin with no significant response (Figure 1d, iv & v). Together, these controls demonstrate the resilience of the formed monolayers to small molecule penetration within the Debye region, enabling MT-facilitated detection without extraneous interference from unbound analytes or other electroactive interferants.

Serotonin-specific MTs were next challenged with increasing concentrations of serotonin in PBS. Chronoamperometric readout showed increasing response waveforms with increasing serotonin concentration from 10 pM to 10 nM, resulting from the additional faradaic contribution of serotonin oxidation (Figure 2a). Serotonin detection was quantified as the change in signal response and reported in Figure 2b. As the concentration of serotonin exceeded 1 nM, the signal response continued to increase but at a noticeably slower rate. Given the sequential nature of the concentration testing, the lower signal response at high concentration may be due to incomplete reduction of the serotonin following previous scans, resulting in receptor-bound oxidized serotonin that cannot contribute to subsequent signal generation.

**Figure 2.**
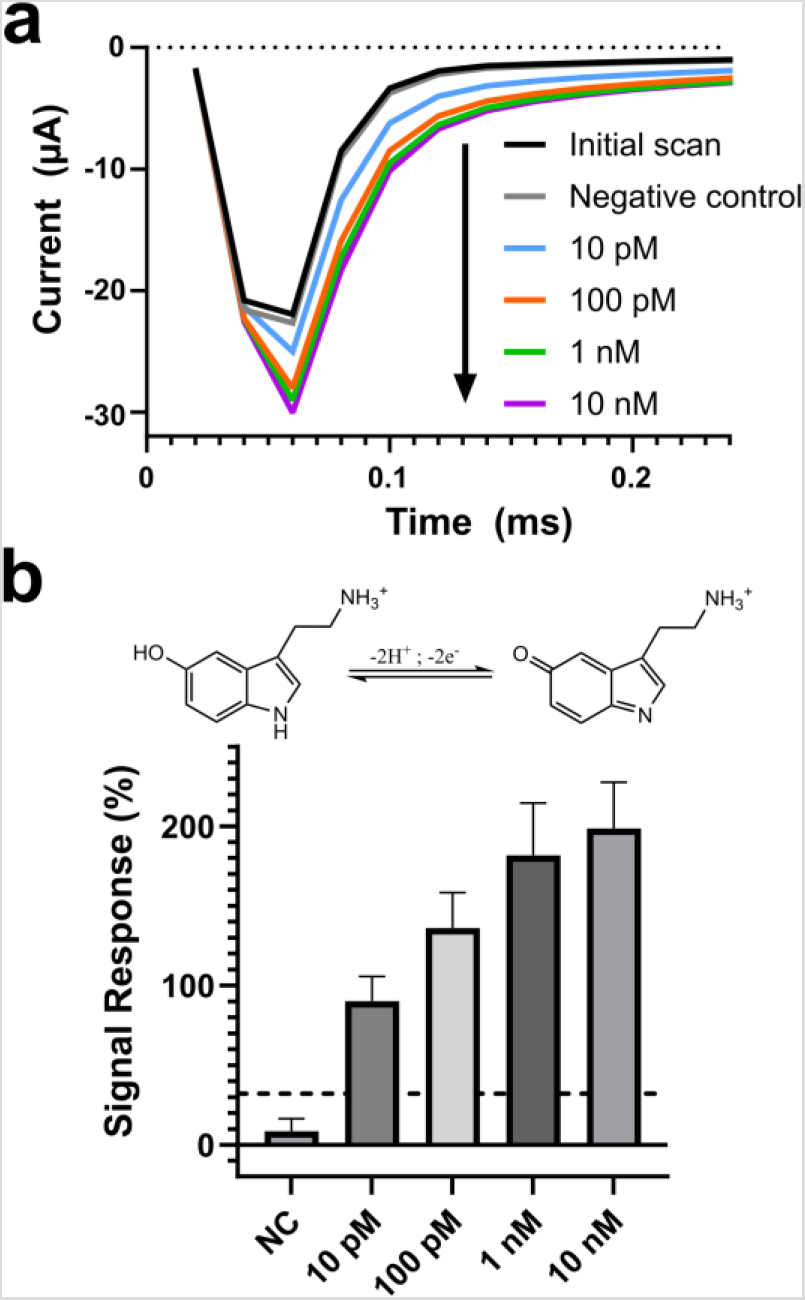
Monolayer transporter serotonin assay. (a) Chronoamperometric response waveforms of serotonin-specific MTs after 30 min incubation in 10 µM dopamine negative control (NC) and various concentrations of serotonin. (b) Obtained signal response of serotonin-specific MTs exposed to serotonin (n = 5, dashed line = detection limit).

To further validate the resilience of the MT monolayers to passive electroactive analyte penetration, serotonin-specific MTs were exposed to high concentration dopamine for an extended time (Figure S5). Over this three-hour period, no significant signal change was observed in the 40-300 µs window; some minor drift was observed in the 20 µs timepoint – most likely due to capacitive changes – but this did not influence the relevant detection time window.

Having established a proof-of-concept MT sensor for serotonin, the approach was next extended to other electroactive small molecules including dopamine, doxorubicin, and epinephrine. Dopamine-specific MTs were challenged with increasing concentrations of dopamine in PBS, which produced a significant response down to 1 pM (Figure 3a). Dopamine signal changes were observed to be larger than those of serotonin, potentially due to the higher observed oxidative conversion seen during the DPV comparison (Figure 1b). This larger signal may be the result of dopamine oxidation falling entirely within the 0-500 mV window. Doxorubicin-specific MTs were similarly challenged with increasing doxorubicin concentration in PBS and exhibited increasing signal response (Figure 3b).

**Figure 3.**
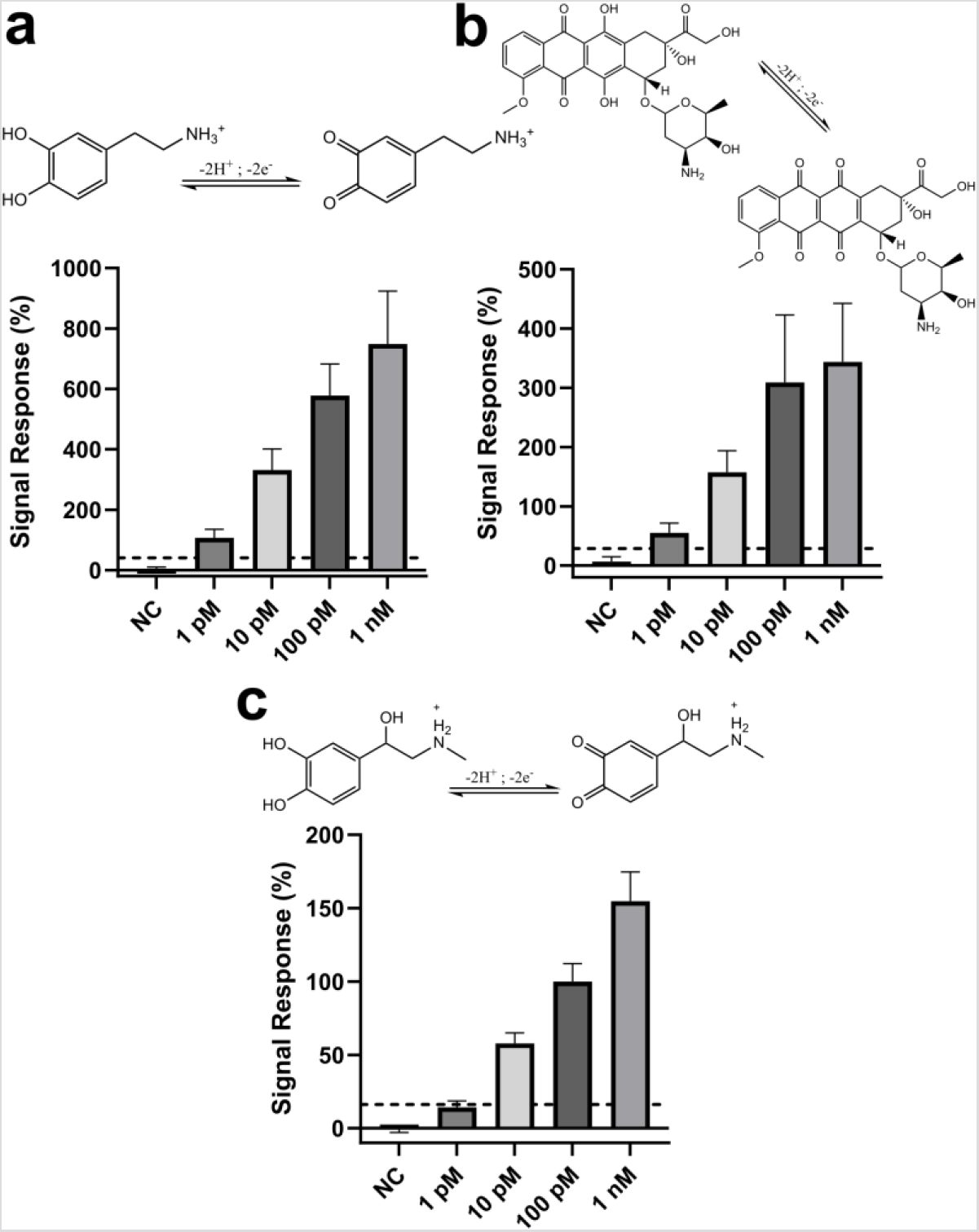
Monolayer transporter analysis of additional molecular targets. MT-based detection of (a) dopamine as well as a negative control (NC, 10 µM serotonin) (b) doxorubicin as well as a negative control (NC, 10 µM cortisol and epinephrine) (c) epinephrine as well as a negative control (NC, 10 µM serotonin) (n = at least 4 for all, dashed line = detection limit).

While the previous serotonin, dopamine, and doxorubicin MTs employed aptamers as the recognition element, antibodies were also tested as viable receptors, using epinephrine as a target. Epinephrine antibody MTs demonstrated a similar upwards trend in signal to the previous aptamer sensors when challenged with increasing epinephrine (Figure 3c). While the observed signal response for epinephrine was below those obtained for the aptamer-based MTs, we believe this is attributable to the larger size antibody receptor being less sterically free to bring the epinephrine within reach of the electrode.

To assess the benefit of the MT approach compared to previous small molecule sensing methods, we explored two alternative mechanisms: direct voltammetry of electroactive analytes on an electrode and direct attachment of an aptamer to the surface of an electrode. Upon exposure of a bare nanostructured gold electrode to a mixture of serotonin, dopamine, epinephrine, and doxorubicin, a concentration of approximately 4 µM was required to achieve a significant response (Figure S6). This is likely due to the lack of localization of small molecules near the electrode surface. After coating a gold nanostructured electrode in a dopamine aptamer and MCH monolayer, and exposing the electrodes to dopamine, a significant response was obtained between 1-10 nM dopamine (Figure S7). While this represents a significant improvement over direct oxidation on bare gold, it is still less sensitive than the MT detection of dopamine. We attribute the improved sensitivity of the MT approach over the surface aptamer approach to the directed movement of the rigid MT towards the surface – in contrast to the random collapse of surface-bound aptamers – as well as the ability of the MT to extend the receptor further into solution and lessen potential steric effects.

To assess the feasibility of employing the MT approach in biological fluids, serotonin specific MTs were challenged with serotonin spiked into a variety of biological fluids. To account for pre-existing, non-degraded endogenous serotonin, sensors were left to equilibrate in the fluid for 30-60 min prior to testing. Baseline measurements were established by taking two, 30 min separated readings and ensuring minimal change was observed; these stability readings were used to establish the limits of detection. Serotonin detection in saliva was achievable down to 100 pM, which was just above the determined limit of detection (Figure 4a). This concentration, as well as the 1 nM treatment, produced relatively low signals compared with higher concentrations (10 nM, 100 nM) that had more pronounced signal changes. This trend was similarly observed with serotonin measurement in sweat (Figure 4b). However, MT sensitivity in sweat was reduced by roughly an order of magnitude, with the 1 nM treatment providing a signal change just above the determined detection limit and the 100 pM treatment showing no significant response.

**Figure 4.**
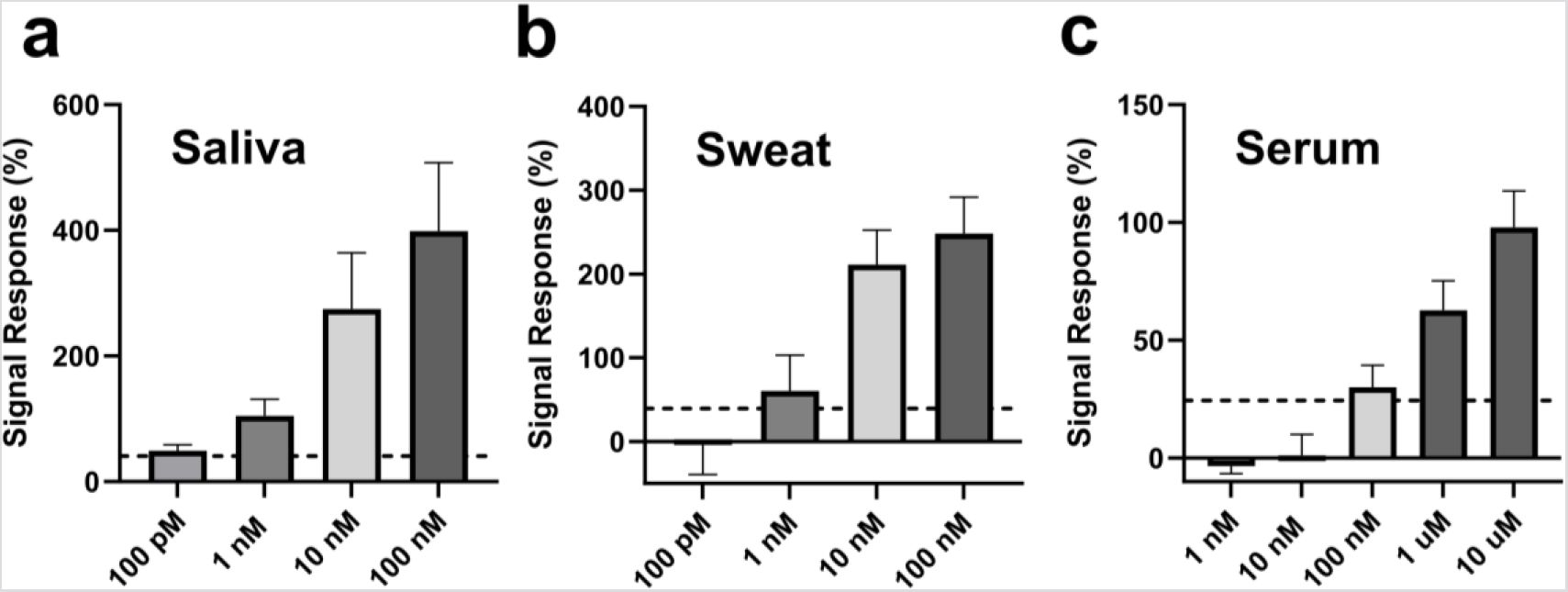
Monolayer Transporter-based Detection in Biological Fluids. Detection of serotonin spiked into human (a) saliva (b) sweat (c) serum (n = at least 3 for all, dashed line = detection limit).

Serum testing of serotonin showed further reduced sensitivity with a limit of detection around 100 nM (Figure 4c), but it is noteworthy that this threshold is below the physiological range of serotonin in blood (∼600-1100 nM) ^19^. We speculate that the reduced sensitivity of MT sensors in serum is due to the combined effects of both the small size of the analyte relative to the receptor and the steric hindrance from electrode-localized serum proteins as well as the endogenous serotonin that may be occupying the MTs. Detection of dopamine in serum was also explored (Figure S8) but proved difficult to accurately assess given the rapid degradation of dopamine in plasma. ^20^

The reusability of the monolayer transporter system was tested by interrogating dopamine-specific monolayer transporter sensors with varying concentrations of dopamine with current sampled every 5 min (Figure 5). The obtained signal responses showed decent agreement with the previous dopamine buffer assay and correlated well with concentration change. It should be noted that rapid sampling of the monolayer transporters led to reduced signal, possibly due to the analyte having insufficient time to completely reduce back to the original form or be replaced by non-oxidized analyte (Figure S9). Thus, a sampling interval of at least 1-2 minutes appears necessary to ensure equilibrium is achieved.

**Figure 5.**
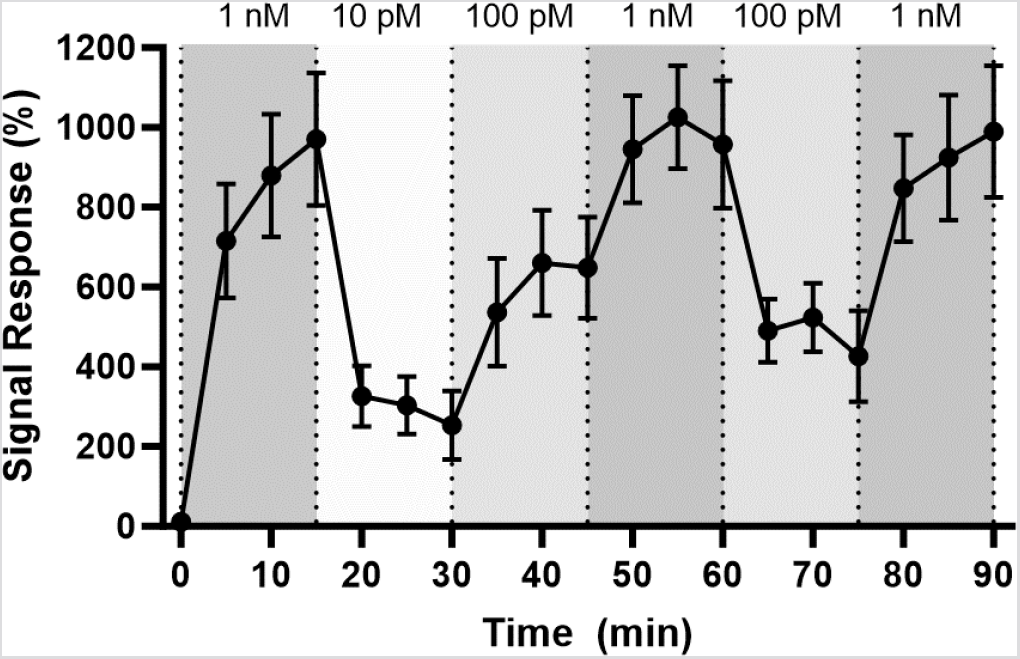
Continuous Monitoring of Dopamine. **D**etection of varying dopamine concentrations in PBS with 5-min sampling (n = 4). Dopamine concentration was changed every 15 min with a brief buffer wash before each sample change.

In summary, we have detailed a new approach to small molecule monitoring that involves directed movement of analyte to the electrode surface for signal generation. This method requires that the small molecule be electroactive under 500 mV (vs. Ag/AgCl) which limits the compatible targets; however, it is likely that this mechanism can be extended to other electroactive analytes (e.g., peptides, proteins) that satisfy the above condition. In fact, we suspect that the large size difference between the receptors used in this study (i.e., aptamers, antibodies) and the small molecule analytes are one reason for the observed sensitivity reduction in biological fluids^21^ – an issue that may be resolved by targeting larger analytes. Alternatively, incorporation of smaller recognition elements (e.g., antibody fragments) may enhance signal resolution for small molecule targets.^22^

The creation of new reagentless sensing techniques for biologically relevant analytes is an important goal for the further development of continuous analysis systems for physiological monitoring.^4^ The monolayer transporter approach described here is compatible with a variety of biological fluids, and future work will focus on the long-term reusability of this sensing system and explore how it can be used to track small molecules *in vivo*.

## Acknowledgements

This work made use of the NUFAB and EPIC facilities of Northwestern University’s NUANCE Center, which has received support from the SHyNE Resource (NSF ECCS-2025633), the IIN, and Northwestern’s MRSEC program (NSF DMR-2308691). This work was completed with startup funding from Northwestern University and a gift from the Chan Zuckerberg Initiative.

