## Supplemental for "Self-Assembled Monolayer Transporters Enable Reagentless Analysis of Small Molecule Analytes"

### Materials.

All DNA sequences were purchased from Integrated DNA Technologies (IDT). The monolayer transporter (MT) sequences are: 5′-SH-MC6-TAC CAG CTA TTG TAT CTA ATA AGA-3′ (anchor strand) and 5′-X-C6-TCT TAT TAG ATA CAA TAG CTG GTA-3’ (receptor strand), where X is either an additional aptamer sequence with a three thymine (TTT) spacer or an amine group (-NH2) for conjugation of the receptor strand to an antibody. Dopamine and serotonin aptamer sequences were obtained from Nakatsuka et al. (2018).[1] The doxorubicin aptamer sequence was obtained from Li et al. (2017).[2] Epinephrine antibody was acquired from Abbexa (UK). Serotonin hydrochloride, dopamine hydrochloride, epinephrine, doxorubicin, 6-mercaptohexanol (MCH), hexaammineruthenium(III) chloride, and tris(2-carboxyethyl)phosphine hydrochloride (TCEP), and human serum were purchased from Millipore Sigma (Canada, USA). Human saliva and sweat were obtained from Lee BioSolutions (USA).

### Receptor Conjugation.

Epinephrine detection involved conjugation of an epinephrine-specific antibody to the 5’ end of the receptor strand. This conjugation was achieved using an oligonucleotide conjugation kit (Abcam, US, ab218260). All other analytes were targeted with aptamers which were easily appended to the end of the receptor strand without the need for additional conjugation.

### Electrochemical Chip Fabrication.

Electrochemical chips were fabricated via photolithography in a class-100 cleanroom. Gold-coated glass wafers with titanium adhesion layers (100nm gold, 5nm titanium) were obtained from Platypus Technologies (USA). Gold electrodes were patterned on the wafers using MICROPOSIT S1813 photoresist, followed by gold/titanium etching. Working electrode apertures (20 μm diameter) were formed using SU-8 photoresist. All photoresists were obtained from Kayaku Advanced Materials (USA).

**Electrochemical Instrumentation.**

Electrochemical analyses were performed using Epsilon Eclipse Potentiostats (BASi, USA). All experiments were conducted using a three-electrode system, with electrochemical chip-localized working electrodes, and external reference (Ag/AgCl) and counter (platinum) electrodes – both of which were also purchased from BASi.

Chronoamperometry measurements were used to monitor detection and involved application of a 500 mV (vs. Ag/AgCl) pulse (with an initial potential of 0 mV) for 50 ms, with a sampling interval of 20 μs.

**Chip Cleaning & Gold Nanostructure Growth.**

Prior to use, cleanroom-fabricated chips were subjected to oxygen plasma for 30s to help remove organic contaminants. Next, chips were cleaned through sequential rinsing with acetone, isopropyl alcohol, and water, followed by drying with nitrogen gas.

Gold nanostructures were grown out of the formed SU-8 apertures through immersion of the chips in 50 mM $HAuCl_4$, followed by direct current potential amperometry with the following 2 step electrodeposition parameters: 1) application of 0 mV (vs. Ag/AgCl) for 100s 2) application of -450 mV (vs. Ag/AgCl) for 10s. The first step in this process enables growth of dendritic gold structures that drastically increase surface area, while the second step produces finer nanostructures to enhance sensitivity. Figure S1 shows a scanning electron microscopy picture of a standard gold nanostructured electrode grown using the above protocol.

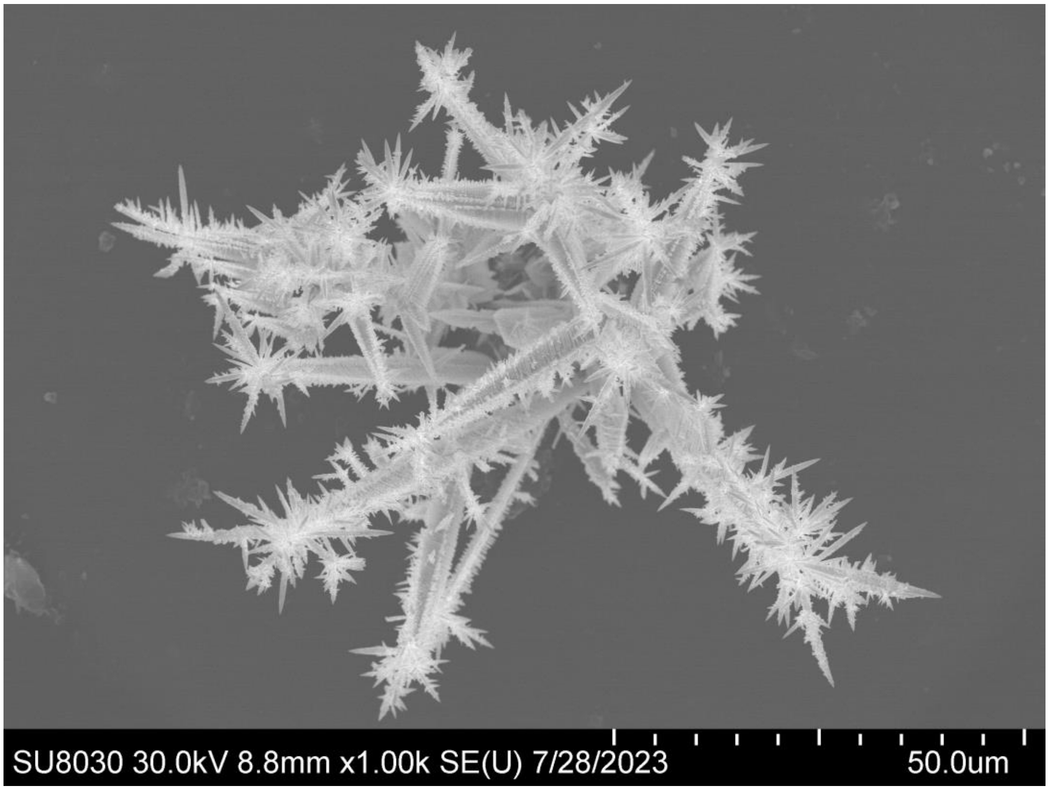


**Figure S1:** Gold nanostructured microelectrode (NME).

Following nanostructure deposition, electrodes were electrochemically cleaned via repeated cyclic voltammetry cycling between 0 and 1.5 V (vs. Ag/AgCl) in 0.1 M sulfuric acid until stable curves were obtained. Figure S2 shows a typical voltammogram for the gold nanostructured electrodes obtained following cleaning. Using the area under the reduction peak and a gold oxide charge transfer value of 390 μC/$cm^2$,[3] an effective surface area of approximately $3.41 \times 10^{-4}$ (±0.31) $cm^2$ (n=21) was determined for the gold nanostructured electrodes employed in this study.

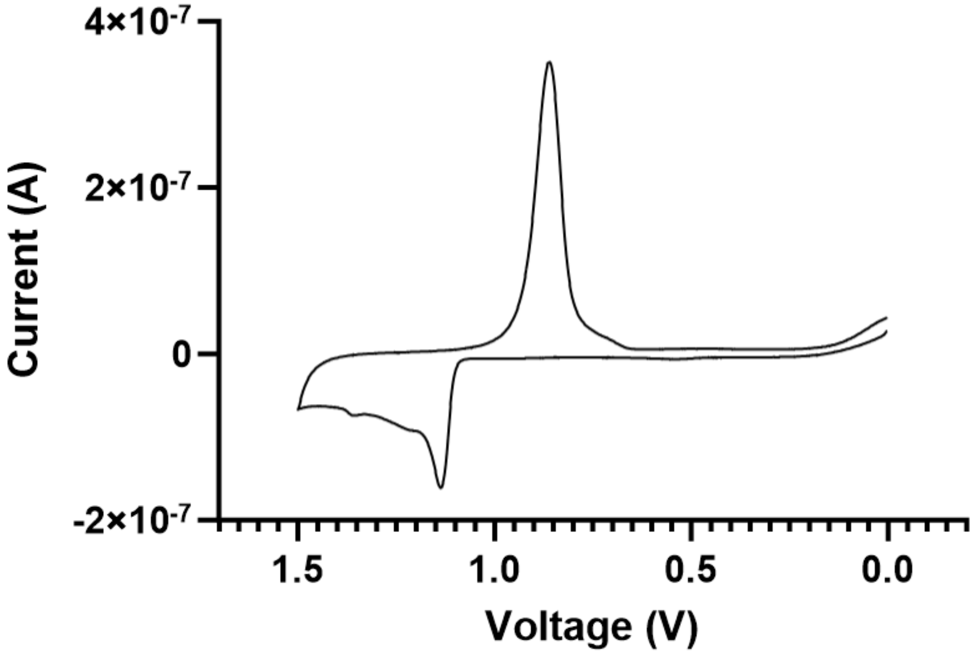


**Figure S2:** Cyclic voltammogram of nanostructured gold electrode in 0.1 M sulfuric acid.

**Electrode Functionalization & Testing.**

MT probes were prepared via hybridization of anchor and receptor strands. Briefly, the anchor strand solution was prepared via dilution of the anchor strand in PBS to a final concentration of 20 μM. In addition, TCEP was added to this solution with a final concentration of 10 mM to help facilitate oligonucleotide disulfide reduction. The anchor solution was left to sit for approximately 10 minutes after mixing and dilution. The receptor strand solution was prepared by dilution of the receptor strand in PBS to a final concentration of 20 μM. Next, anchor and receptor strand solutions were mixed in a 1:1 ratio for a final overall MT probe concentration of 10 μM. This mixed solution was heated for 10 minutes at $95^0$C, then left to cool to room temperature for approximately 20 minutes. After cooling, the probe solution was ready for deposition. It should be noted that this protocol was employed for all MT tests in PBS. For testing in biological fluids, a slightly modified protocol using a final probe concentration of 1 μM was used as this order of magnitude decrease in starting probe produced higher signal resolution. We speculate the lower probe density improved signal resolution in biological fluids by providing more lateral space for the DNA constructs to fall to the surface, possibly counteracting blocking caused by protein adsorption.[4] It should also be noted that the $95^0$C heating step was only performed for aptamer-based recognition elements. For the antibody receptor (epinephrine), the anchor strand was heated to $95^0$C, cooled slightly, and then combined with the receptor strand.

Following electrochemical cleaning of the working electrodes, the final probe solution was deposited onto the nanostructured electrodes and left to incubate overnight (12-14h) at room temperature in a humidity chamber. After probe incubation, the probe solution was gently removed from the electrodes and replaced with a solution of 1 mM MCH. The electrodes were left to incubate in MCH for 2 hours at room temperature in a humidity chamber. Following MCH incubation, the chips were thoroughly rinsed with PBS via repeated buffer swapping (at least five times with 3-minute equilibration periods). Finally, electrodes were left in PBS for 10-20 minutes to help condition the monolayer to ensure stability. Prior to any analyte measurement, two chronoamperometry measurements were taken 30 minutes apart and compared to ensure stable readout. If drift was observed, electrodes were incubated for another 30 minutes in 10 mM MCH and the process was repeated. If after two 30-minute 10 mM incubations the chips were not stable, then these chips were discarded. Most chips were stable after the 1 mM MCH incubation or an additional 10 mM MCH incubation; a few of the tested chips required the second 10 mM MCH backfill, and a few were discarded due to instability issues.

Following the buffer stability test, electrodes were incubated in a high concentration negative control (10 μM of another non-specific electroactive analyte) for all PBS measurements prior to analyte testing to ensure monolayer integrity. Thus, all PBS measurements underwent two controls: stability in buffer and stability with negative control. For biological fluid testing, following the initial buffer stability test, PBS was replaced with the biological fluid and chips were left to equilibrate in the solution for 30-60 minutes. Following this equilibration period, a 30-minute biological fluid stability test – like the buffer stability test – was performed to ensure stability of the sensors in the fluid. This test was used as the negative control for the biological fluid given the high amount of electroactive material in the solution.

When the sensors had passed both the buffer stability and negative control tests (for PBS data) or both the buffer stability and biological fluid stability tests (for biological fluid data), the sensors were cleared for analyte interrogation. The sensors were incubated for 30 minutes in various concentrations of the analyte spiked into either PBS or a biological fluid. Chronoamperometry measurements were performed following the 30-minute incubation and compared with the baseline measurement (i.e., stability test).

For the continuous dopamine test, dopamine-specific MT sensors were prepared and incubated in varying concentrations of dopamine in PBS after ensuring electrode stability. The concentration of dopamine was changed every 15 min, with a brief buffer rinse before each replacement. Chronoamperometric signals were measured every 5 min for a total of 90 min. An additional measurement was performed at 80.5 min to assess the effect of rapid measurements on MT readout.

**Electrode Probe Density Measurement.**

Given the apparent reliance of the MT approach on probe density, the concentration of MT probes on the electrode was determined using ruthenium hexamine-assisted chronocoulometry.[5] Electrodes were functionalized according to the above reported protocol with 10 μM serotonin MT probe overnight, followed by 2 hours in 1 mM MCH and the necessary buffer washes. Next, chronocoulometric measurements were taken by applying a 250 ms pulse from 200 to -500 mV (vs. Ag/AgCl) to the electrode in 0.1X PBS to generate a plot of charge vs. sqrt(time) (Figure S3). This test was repeated on the same electrode after exposure to nitrogen-purged 100 μM $[Ru(NH_3)_6]^{3+}$ which is expected to adhere to DNA and increase charge transfer. Using the difference in charge, and the previously determined electrode surface area, the probe density was determined to be approximately $4.3 \pm 0.74 \times 10^{12}$ probes/cm$^2$ or 1 probe per 23 nm$^2$.

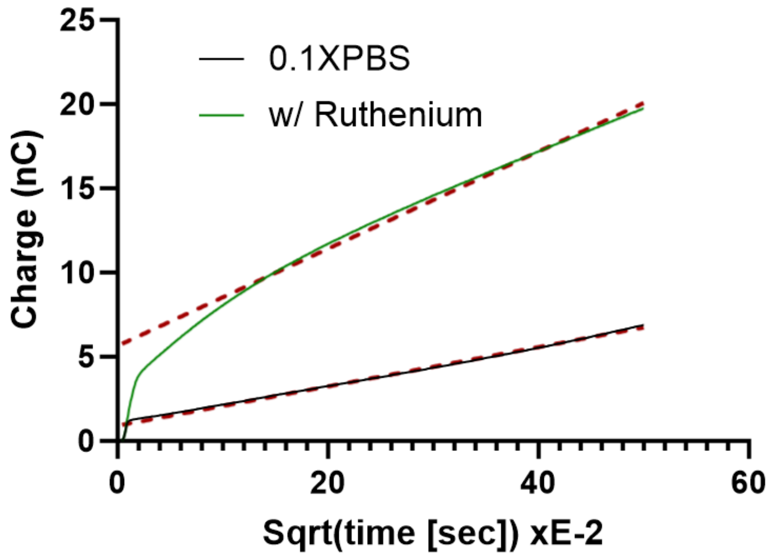


**Figure S3:** Chronocoulometry curves for serotonin MT-functionalized sensor in the presence and absence of 100 μM $[Ru(NH_3)_6]^{3+}$ in 0.1X PBS.

**Data Analysis**

Chronoamperometry data was analyzed and compared by simple comparison of current at a selected time point. In this case, the 140 μs point was selected for all experiments as it consistently showed the largest difference among treatments. Signal responses were reported by calculating the percentage change in the current at 140 μs relative to an initial baseline measurement. For example, a 100% sensor response represents a 100% increase in current at the 140 μs timepoint compared to the baseline. This baseline standardization was employed to help correct for minor differences among electrodes (e.g., surface area and probe number).

**Detection of Serotonin with NME and Flat Gold Electrodes.**

While NMEs were employed for all experiments performed in this study, we also sought to explore whether MT-based detection was possible using standard flat gold electrodes. For this test, gold electrodes were fabricated in the cleanroom with working electrode apertures that had surface areas approximately equal to the previously calculated effective NME surface area ($3.5 \times 10^{-4}$ $cm^2$). Next, both electrode types were functionalized with serotonin-specific MT monolayers and tested to ensure stability in buffer. Finally, both electrode types were challenged with 1 nM serotonin in PBS for 30 min. While both electrode types saw a noticeable signal change upon serotonin addition (Figure S4), the NME signal was drastically larger than that of the flat gold electrodes.

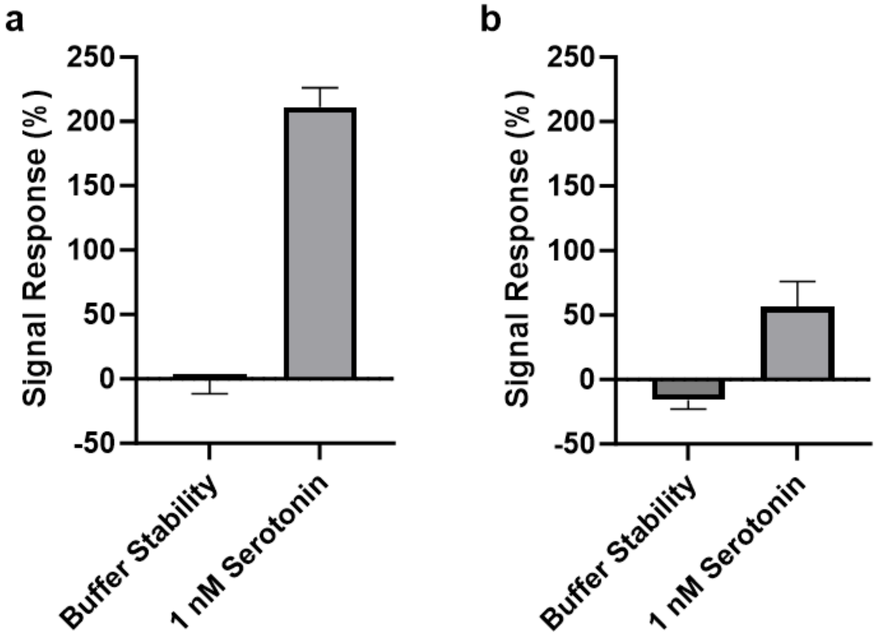


**Figure S4:** Detection of 1 nM serotonin in PBS using serotonin-specific MTs on both (a) NME electrodes (b) flat gold electrodes of approximately equal surface area (n=3 for all).

**Stability of Serotonin MT Monolayers.**

Stability of serotonin-specific MTs was assessed over an extended period of time through exposure to a high concentration negative target (10μM Dopamine). During the 3-hour period, no significant drift was observed in the 40-300 μs window used for detection (Figure S5).

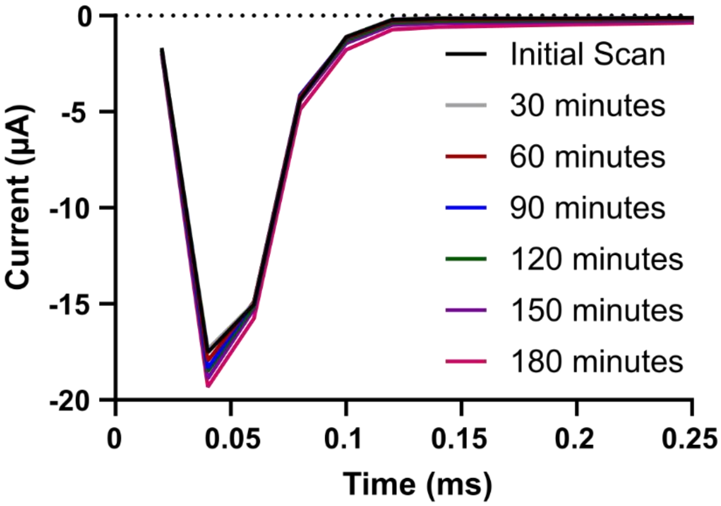


**Figure S5:** Chronoamperometric response waveforms of serotonin-specific MTs exposed to 10 µM dopamine for 3 hours with measurements taken every 30 min.

**Detection of Electroactive Species using Direct Voltammetry.**

The ability of a bare gold nanostructured electrode to detect electroactive small molecules through direct voltammetry was assessed. For this, solutions consisting of a mixture of the small molecules employed in this study (serotonin, dopamine, epinephrine, doxorubicin) were used to challenge the electrodes. Upon exposure to these solutions, the sensors produced a significant signal only at or above 4 uM (Figure S6).

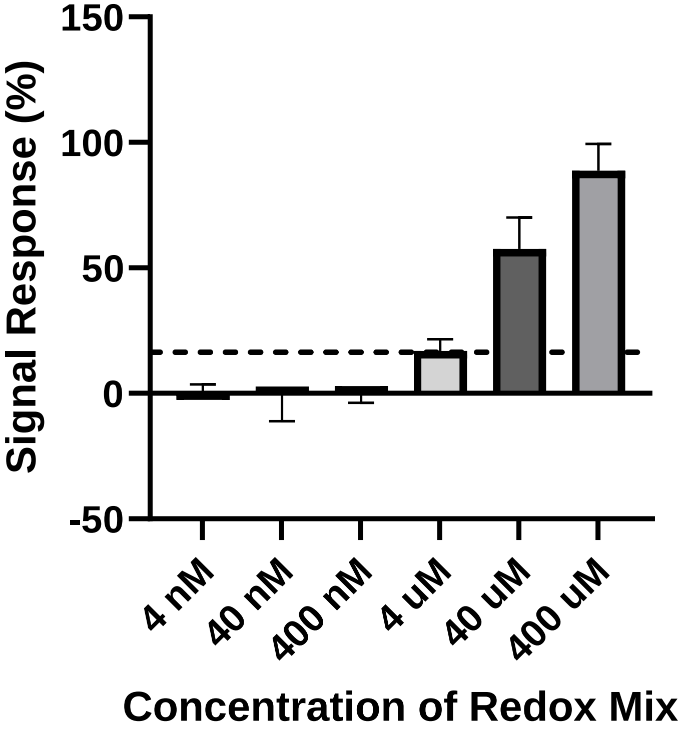


**Figure S6:** Detection of a redox mixture composed of equal parts serotonin, dopamine, epinephrine, and doxorubicin exposed to a bare gold nanostructured electrode. (n = at least 6)

**Detection of Dopamine using Direct Aptamer Monolayers.**

The ability of gold nanostructured electrodes modified with aptamers directly on the surface was assessed. Sensors were prepared by attaching dopamine aptamers directly to the electrode surface along with MCH according to the same deposition protocol as used for the MT approach. Upon being challenged with increasing dopamine concentrations, the sensors produced a significant response only after 1-10 nM of dopamine was present (Figure S7).

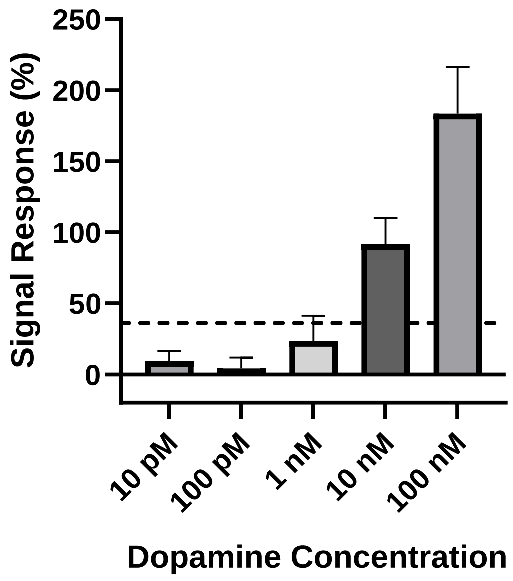


**Figure S7:** Detection of dopamine using a dopamine aptamer-modified gold nanostructured electrode. (n = at least 4).

**Detection of Dopamine in Human Serum.**

Dopamine detection with MTs was also explored in biological fluids. However, given the rapid reported half-life of dopamine in plasma (~2 min), it proved difficult to accurately quantify the dopamine after 30 min incubation periods. Regardless, high initial spiked concentrations of dopamine showed some signal response (Figure S6).

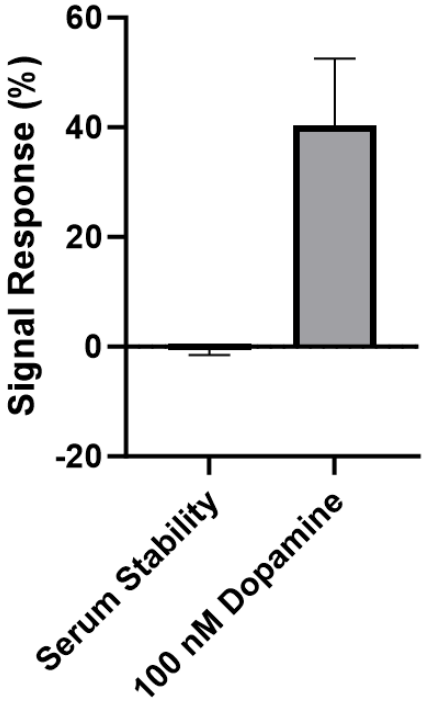


**Figure S8:** Detection of dopamine spiked into serum (n=4).

**Rapid Sampling of Dopamine Concentration.**

Continuous dopamine detection with MTs was performed in PBS with varying concentrations of dopamine (Figure 5). As part of this study, measurements were also performed 30s after the 80 min timepoint (1 nM dopamine present) – instead of the typical 5-minute sampling interval – to test the effects of rapid sampling. This datapoint (shown as the blue circle in Figure S9) showed reduced signal response compared to the 80-, 85-, and 90-min measurements, likely due to insufficient time for full reduction of the dopamine back to its base form or replacement of the oxidized dopamine with the unoxidized form. Thus, it appears that a sampling interval of at least 1-2 minutes is necessary to allow for full equilibration and ensure accurate readings of bulk concentration.

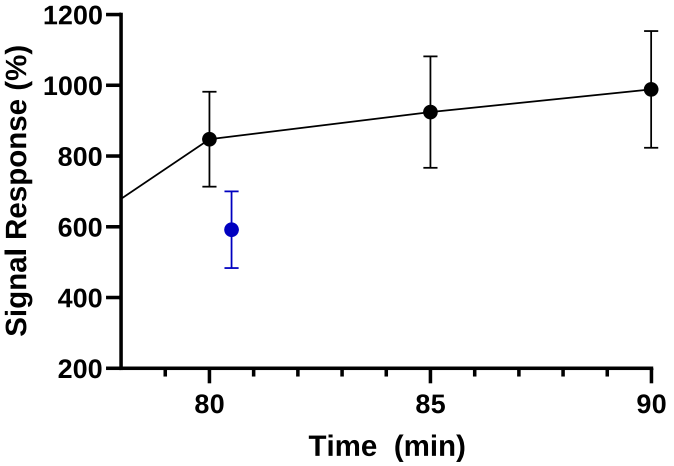


**Figure S9:** Continuous sampling of 1 nM dopamine in PBS (n=4).

### References (Supporting Information):
